# RodZ promotes MreB polymer formation and curvature localization to determine the cylindrical uniformity of *E. coli* shape

**DOI:** 10.1101/226290

**Authors:** Randy M. Morgenstein, Benjamin P. Bratton, Joshua W. Shaevitz, Zemer Gitai

## Abstract

Cell shape in bacteria is determined by the cell wall, which is synthesized by a variety of proteins whose actions are coordinated by the actin-like MreB protein. MreB uses local geometric cues of envelope curvature to avoid the cell poles and localize to specific regions of the cell body. However, it remains unclear whether MreB’s curvature preference is regulated by additional factors, and which features of MreB are essential for specific aspects of rod shape growth, such as cylindrical uniformity. Here we show that in addition to its previously-described role in mediating MreB motion, RodZ also modulates MreB polymer number and curvature preference. MreB polymer number and curvature localization can be regulated independently. Quantitative 3D measurements and a series of mutant strains show that among various properties of MreB, polymer number, total length of MreB polymers, and MreB curvature preference are the key determinants of cylindrical uniformity, a measure of the variability in radius within a single cell. Changes in the values of these parameters are highly predictive of the resulting changes in cell shape (r^2^=0.93). Our data suggest a model for rod shape in which RodZ promotes the assembly of multiple long MreB polymers that ensure the growth of a uniform cylinder.

## Introduction

Understanding how cells encode the ability to robustly determine their own shapes remains one of the central mysteries of cell biology. In bacteria, the peptidoglycan cell wall (PG) forms a rigid structure whose shape dictates the shape of the cell. When purified, the extracted PG maintains the cell’s original shape and loss of PG causes cells to lose their shapes, e.g., rod-shaped bacteria becoming round^1-3^. These cells can then reestablish cell shape *de novo* once cell wall synthesis is restored^2^. In this work, we focus on the gram-negative rod-shaped bacterium *Escherichia coli*. Despite their relative simplicity, there are multiple parameters that describe a population of rod like cells including: a straight cylindrical axis with high uniformity of the diameter (which we term cylindrical uniformity), center-line straightness (bent vs straight rod), the distribution of cell widths and lengths within the population, and the geometry of the poles. Previously we studied the determinants of *E. coli* straightening and population-average width, but the determinants of single-cell cylindrical uniformity remain unclear^2,4,5^.

Computational modeling has shown that rod shape can be established and maintained by directing the insertion of new cell wall material in an organized fashion^6,7^. If cell wall synthesis occurs randomly throughout the cell, the cell becomes unstable and defects become amplified, leading to a loss of rod shape. In *E. coli*, the bacterial actin homolog MreB organizes cell wall insertion by localizing to regions of the cell with particular geometric curvatures and recruiting cell wall enzymes to direct growth to those sites^2,4^. However, it remains unknown whether accessory proteins regulate MreB assembly or curvature-preference. Furthermore, it remains unclear which combination of MreB’s properties are necessary to direct specific aspects of rod shape formation.

In *E. coli*, cell wall insertion is localized to the main body of the cell, with no growth at the poles where the cell wall remains inert^8^. The lack of cell wall insertion at the poles can be explained by geometric exclusion of MreB, which directs the locations of cell wall insertion^4^. Because the cell poles are much more curved than the rest of the cell and MreB is excluded from regions with such curvature, sensing cell curvature is a powerful way for MreB to recognize and avoid the poles. However, the scale of cellular curvature is much larger than that of a single protein, making it difficult for monomeric proteins to detect the difference between polar and mid-cell geometry^6,9^. To overcome this problem, MreB has been proposed to form cellular-scale polymers, whose assembly has been observed both *in vivo* and *in vitro*^10-13^.

MreB polymers serve multiple roles in cell shape determination. First, elongated polymers can coordinate the activity of multiple cell-wall modulating enzymes to produce twisting cylindrical growth^14^. Second, the orientation of MreB polymers relative to the cell axis helps determine the average cell width of the population^5^. Lastly, MreB polymers allow these nano-scale proteins to form micron-scaled structures that can sense membrane curvature differences between the poles and main cell body. More specifically, *in vivo* localization studies show that MreB is depleted from the poles and is enriched at areas of low or negative mean curvature^4^. *In vitro*, MreB filaments can bend a membrane vesicle and molecular dynamics simulations suggest that MreB polymers have an intrinsic bend^11,15^. We previously analyzed the properties of MreB that correlate with the population-level average width of the cell and found that MreB polymer angle correlates with average cell width^5^. However, previous studies have not examined the MreB properties coupled to the ability to form an elongated cylindrical rod-like cell in the first place.

Several toxins have been proposed to target MreB under conditions of stress^16-18^, but it remains unclear whether MreB assembly or curvature localization are normally regulated *in E. coli*. RodZ is a transmembrane protein (Fig. 1A) that is co-conserved with MreB^19^ and is one of the few proteins that definitively binds MreB, *in vitro* by co-crystallization and *in vivo* by bimolecular fluorescence complementation (BiFC)^1,20^. We previously showed that RodZ functions downstream of MreB as an adaptor that causes MreB to rotate around the cell circumference; RodZ couples cytoplasmic MreB to the periplasmic activity of cell wall synthesis^1^. This dynamic rotation promotes robust rod shape in the presence of cell wall stress.

**Figure 1.**
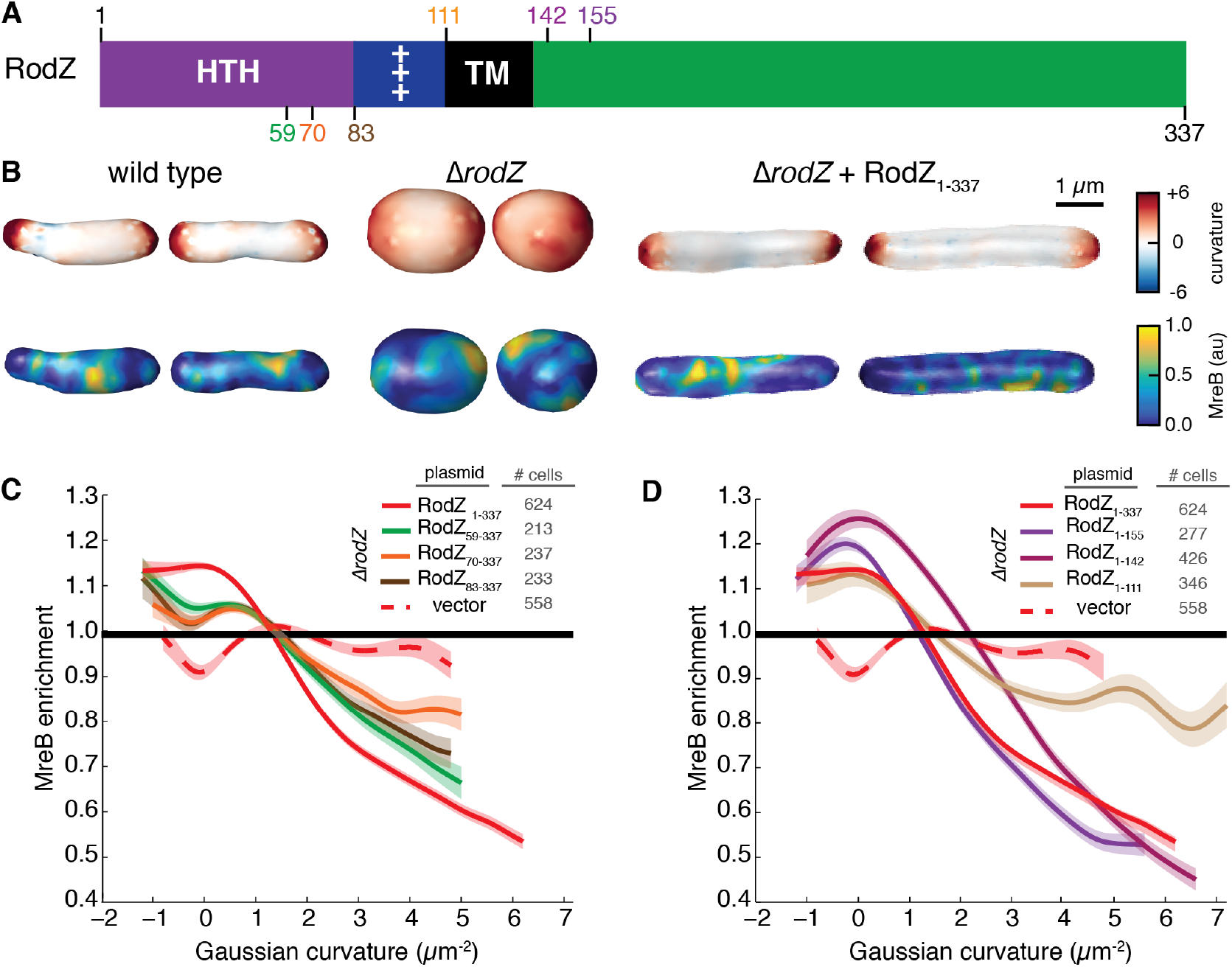
The cytoplasmic domain of RodZ is necessary for MreB curvature localization. A) Schematic of RodZ’s domain structure with location of truncations noted. B) 3D images of WT, Δ*rodZ*, and *rodZ* complemented cells with Guassian curvature and MreB fluorescence intensity represented. C) MreB enrichment plot of RodZ cytoplasmic truncations. D) MreB enrichment plot of RodZ periplasmic truncations. Shaded area indicates ± 1 standard error of the mean, and dotted lines indicate strains deleted of *rodZ*. The curve for each strain is a cubic smoothing spline and is truncated using a probability threshold for extreme curvatures of p>5x10^−3^ (Fig. S2A). Because the shape of each strain is different, the ranges of curvatures plotted for each strain are different.

Here we show that in addition to its role in promoting MreB rotation, RodZ is also a key regulator of MreB assembly and curvature preference. These functions require both the cytoplasmic and periplasmic domains of RodZ, indicating that RodZ functions as a key hub to integrate information across the inner membrane and organize cell shape. Using three-dimensional (3D) imaging and a combination of *mreB* and *rodZ* mutants, we go on to explore which of the many properties of MreB are important for cylindrical uniformity. We find that curvature preference is necessary but not sufficient to grow cylindrically uniform cells, while a combination of MreB polymer number, total polymer length, and curvature preference accurately predict changes in cylindrical uniformity.

## Results

### RodZ is required for MreB curvature localization

We recently showed that the transmembrane protein RodZ interacts with both MreB and the cell wall synthesis machinery to couple MreB rotation to cell wall synthesis^1^. RodZ is necessary for MreB rotation and specific point mutations in *mreB* can roughly restore rod-like shape without restoring MreB rotation^1^. While these data indicated that MreB rotation is not necessary for rod shape, the resulting cells had an irregular morphology distinct from wild-type (WT) cells, suggesting that RodZ could play an important role in the cylindrical uniformity of cell shape independently of its role in MreB rotation. Consequently, we examined the role of RodZ in controlling the biophysical properties of MreB that are thought to be important for shape determination, like curvature preference.

To quantify the effect of RodZ on MreB curvature preference we measured the 3D cell shape and curvature enrichment of MreB in a strain expressing MreB-GFP^sw^ (internal msGFP sandwich fusion) as the sole copy of MreB (Fig. 1B). We previously showed that this fusion fully complements the shape of WT *E. coli* under a wide range of conditions^5^ and all mutants described below were generated in this strain background. Generating 3D cell-shape reconstructions with roughly 50 nm precision from the raw fluorescence images allowed us to calculate the Gaussian curvature, which is the product of the two principal curvatures, at every location on the 3D surface of the cell^21^. These two principal curvatures can only be measured in 3D. Previously we reported MreB’s curvature preference as a function of mean curvature, the average of the two principal curvatures. Mean curvature is sensitive to global properties such as cell size, whereas Gaussian curvature enables us to focus on the local curvature geometry, which is particularly important in irregularly-shaped cells such as Δ*rodZ* mutants. MreB was enriched at Gaussian curvatures near zero and strongly depleted from regions with positive curvature (Fig 1CD). Cell poles have a positive Gaussian curvature since each of the principal curvatures at the pole have the same sign, while cylinders have a Gaussian curvature of zero owing to the lack of curvature along the cell axis. Thus, MreB’s curvature preference nicely parallels the pattern of *E. coli* growth during elongation, localizing to cylindrical regions and avoiding the poles.

Interestingly, we found that deletion of *rodZ* strongly reduced the curvature preference of MreB (Fig. 1CD). In Δ*rodZ* cells, MreB is no longer enriched near zero Gaussian curvature or excluded from the poles. The shape of Δ*rodZ* cells can be complemented by expressing full-length RodZ from a plasmid (RodZ_1-337_). RodZ_1-337_ also restores both the depletion of MreB from regions of positive Gaussian curvature and the enrichment of MreB in regions of negative Gaussian curvature. These cells lacked the peak in MreB enrichment near zero Gaussian curvature noticeable in WT cells (Fig. 2A). The cells we studied do not display much negative Gaussian curvature so the fact that cell shape was similar in WT and RodZ_1-337_ led us to define the important features of WT-like curvature preference as enriched at negative and slightly positive Gaussian curvatures and depleted at strongly positive Gaussian curvatures.

**Figure 2.**
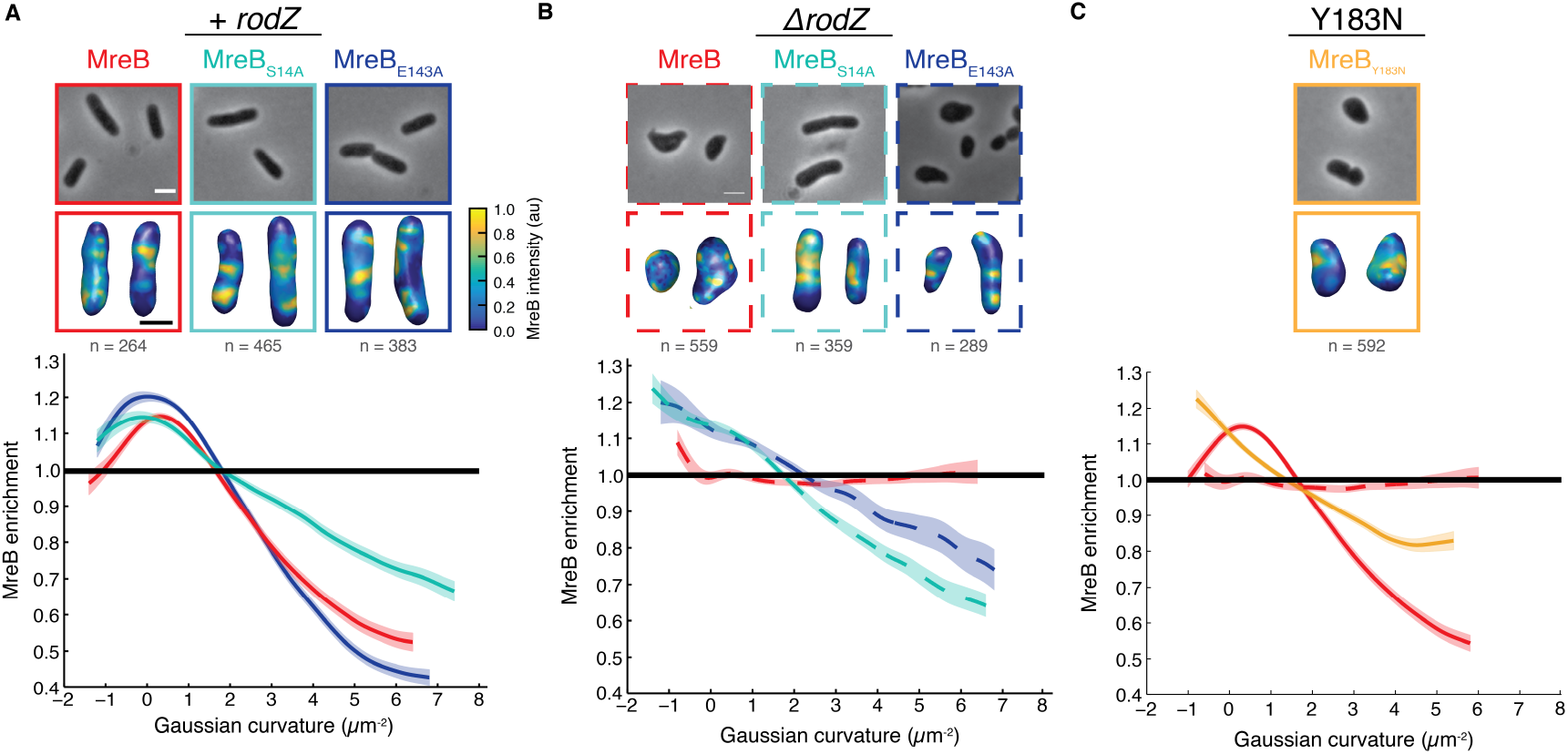
MreB curvature localization is necessary but not sufficient for rod shape. A) MreB enrichment curves of MreB point mutants with RodZ present. B) MreB enrichment curves of MreB point mutants in a *rodZ* deletion. C) MreB enrichment curve of MreB_Y183N_. Top images are 2D cells and bottom images are 3D cells with MreB shown according to the color intensity scale in A. The number of independent cells that contributed to the enrichment plots is indicated in gray. Shaded areas of the curves indicate ± 1 standard error of the mean and dotted lines indicate strains deleted of *rodZ*. The curve for each strain is a cubic smoothing spline and is truncated using a probability threshold for extreme curvatures of p>5x10^−3^ (Fig. S3). Because the shape of each strain is different, the ranges of curvatures plotted for each strain are different. White scale bar for all phase images is 2 μm and the black scale bar for all 3D reconstructions is 1 μm.

Because Δ*rodZ* disrupts both curvature localization and rod shape, we sought to determine if it is possible for MreB to sense curvature in other round cells. We thus grew WT cells in sub-lethal concentrations of the PBP2-inhibiting drug, mecillinam. PBP2 acts downstream of MreB, such that we predicted that PBP2 inhibition would round cells without disrupting MreB’s curvature preference. Additionally, cells were imaged in the absence of mecillinam. We found that in non-rod shaped mecillinam-treated cells, MreB maintained a preference for Gaussian curvatures near and below zero and an avoidance of positive Gaussian curvature (Fig. S1A-C). Because both mecillinam-treated cells and Δ*rodZ* cells lack rod-like shape but only Δ*rodZ* cells lack geometrically localized MreB, the lack of MreB enrichment in Δ*rodZ* must not be a failure in our 3D analysis. These data show that RodZ specifically promotes MreB’s curvature localization in a manner that is not merely secondary to its role in cell shape determination.

### The cytoplasmic domain of RodZ modulates MreB curvature localization

RodZ is a transmembrane protein with a large periplasmic domain and a smaller cytoplasmic domain (Fig. 1A). We hypothesized that these two domains of RodZ could play distinct roles, with the periplasmic domain binding the PG synthesis machinery to promote MreB rotation, and the cytoplasmic domain binding MreB to promote its curvature preference. In order to determine how RodZ regulates MreB curvature localization, we thus examined MreB curvature enrichment in RodZ truncations from both its periplasmic and cytoplasmic termini.

Consistent with our hypothesis, we found that the periplasmic domain plays little role in modulating MreB’s curvature preference (Fig. 1D) even though it is necessary for cell shape^1,12,22^. For example, the curvature localization of RodZ_1-155_ is largely indistinguishable from that of RodZ_1-337_, and RodZ_1-142_ retains the overall WT-like pattern of localization. Even after deleting the entire periplasmic domain along with the transmembrane domain (RodZ_1-111_), there is still a noticeable enrichment around zero Gaussian curvature and a steady decline in enrichment as the Gaussian curvature becomes more positive. This indicates that even when RodZ is not in the membrane, its cytoplasmic domain can influence MreB’s curvature preference. Unlike the membrane-bound periplasmic truncations (RodZ_1-155_ and RodZ_1-142_), a small truncation in the RodZ cytoplasmic domain (RodZ_59-337_) shows a clear change in MreB curvature preference with both a decreased enrichment near zero and less of a depletion from regions with positive Gaussian curvature (Fig. 1C) and a concurrent loss of cell shape (Fig. S2C-E). Additional truncations (RodZ_70-337_) and deletion of the entire helix-turn-helix motif (RodZ_83-337_) show a further dampening of the curvature enrichment profile. While they fail to complement MreB curvature localization, cytoplasmic truncations do generate different cell shapes, suggesting that these truncations are being stably expressed. For all of the Gaussian curvature preference measurements reported in this study, we take into account the distributions of curvatures observed such that changes in curvature preference are not due to changes in the available curvatures in the cell (Fig. S1-S3).

### Rod-shaped cells have WT-like MreB localization but WT-like MreB localization is not sufficient for proper cell shape

Deleting *rodZ* results in a loss of cylindrical uniformity that can be suppressed by a point mutant in *mreB* (MreB_S14A_) without restoring MreB rotation^1,23^. Because RodZ is needed for MreB’s proper curvature localization, we determined whether MreB_S14A_ can also suppress the loss of MreB’s curvature enrichment in the absence of *rodZ*. In contrast to its effects on MreB rotation, the curvature enrichment profile of MreB_S14A_ was restored to a WT-like profile in Δ*rodZ* cells (enriched near zero, a sharp decline toward positive curvatures, and depleted at strongly positive Gaussian curvature, Fig. 2AB).

To test whether the correlation between shape and MreB localization observed for MreB_S14A_ Δ*rodZ* is generalizable, we examined additional MreB point mutations that were originally identified as resistant to an MreB targeting drug, A22, and previously characterized^5^. We confirmed that the steady state levels of MreB are not dramatically affected by these point mutations in the presence or absence of *rodZ* (Fig. S4). MreB_E143A_ had little to no effect on MreB localization in the presence of RodZ while MreB_E143A_ Δ*rodZ* restored MreB localization to a WT-like profile (Fig. 2AB). Despite their qualitatively similar MreB curvature enrichment profiles, MreB_S14A_ Δ*rodZ* formed rods that closely resembled WT cells while MreB_E143A_ Δ*rodZ* cells were more irregular (Fig. 2B, Table S4). Analysis of another point mutant, MreB_Y103N_, reinforced the conclusion that proper curvature localization is insufficient for proper rod shape (Fig. S5). MreB_Y103N_ failed to form rods, in the presence of RodZ, despite displaying a WT-like pattern of MreB localization (Fig. 2C). Thus, all the rod-shaped cells we analyzed have geometrically-localized MreB, but not all cells with geometrically-localized MreB form rods.

### RodZ modulates the number of MreB polymers

Given that RodZ promotes both MreB rotation and curvature preference, we sought to determine if there are additional properties of MreB under RodZ control. Specifically, our 3D analysis enabled us to determine MreB polymer length, number, angle with respect to the long cell axis, and fraction of membrane-associated protein. Comparing cells with and without RodZ, we observed that there were not substantial changes in polymer length (Fig. 3) or average polymer angle (Fig. S4A). There was a statistically-significant change in the fraction of MreB associated with the cell periphery (Fig. S4D), but this change was small (~5%) and also observed in mecillinam-rounded cells, such that it does not appear to be a major component of RodZ’s influence on MreB. We note that our forward convolution method^5^ can accurately identify the presence of polymers regardless of their length and can determine the length of polymers greater than 200 nm. Thus, we can calculate the average MreB polymer length for the ~60% of polymers that are longer than 200nm. When these MreB polymers are measured in strains with or without RodZ, there is not a significant difference in the average MreB polymer length (WT= 515 ± 15 nm, *ΔrodZ=* 509 ± 18 nm) (Fig. 3B), nor in the fraction of detected structures >200nm (57%, 59%).

**Figure 3.**
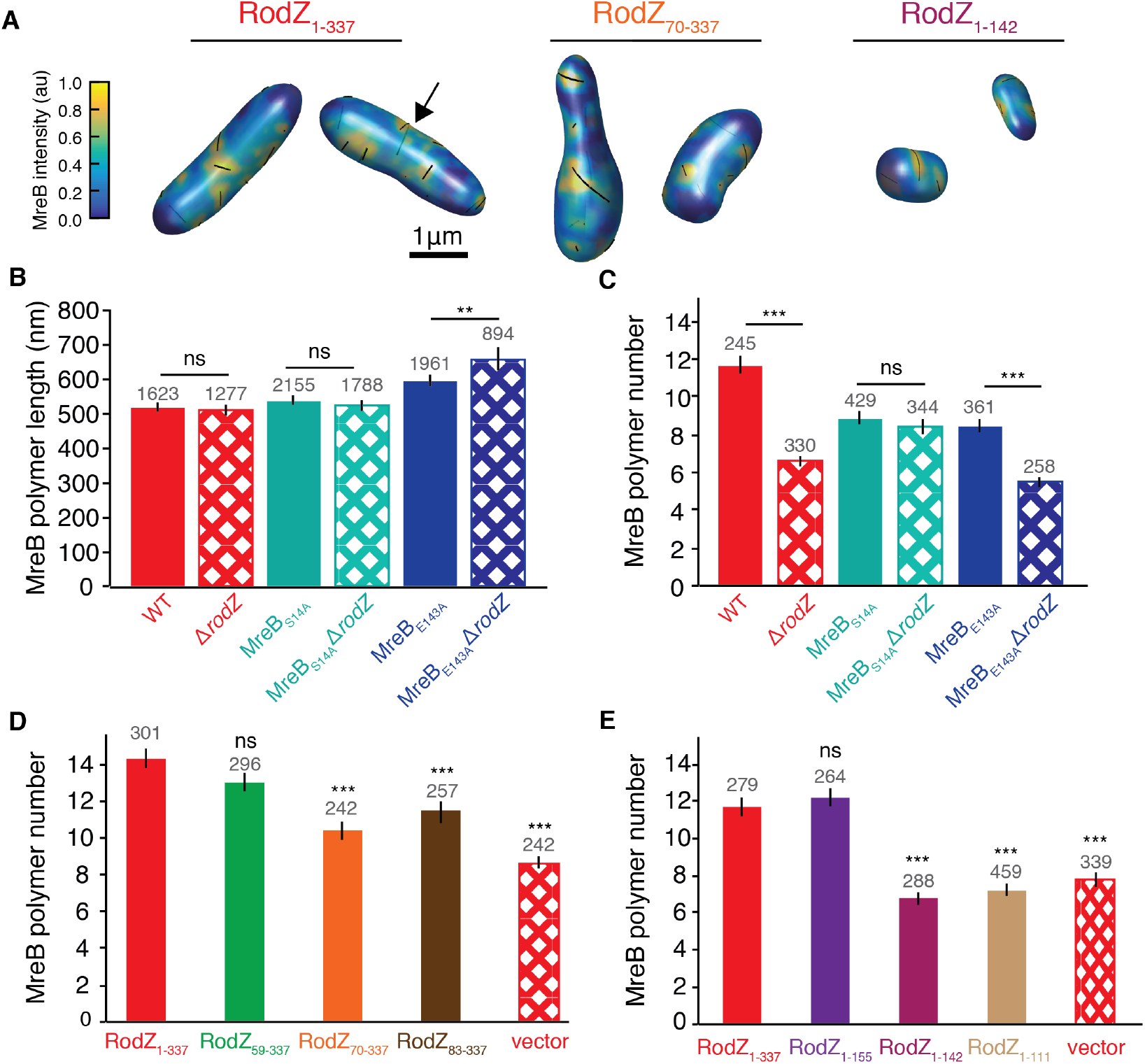
RodZ acts as an MreB assembly factor. A) Semi-transparent 3D renderings of cells with full-length or truncated RodZ as indicated. MreB polymers are indicated with black lines and may be present on the back of the cell where they appear less vividly black (see example at arrow). B) The average MreB polymer length (>200nm) per cell in different MreB point mutants in the presence and absence of *rodZ*. C) The average MreB polymer number per cell in different MreB point mutants in the presence and absence of *rodZ*. B-C) P-value comparisons are made between strains with similar MreB point mutations. For additional cross-comparisons see Table S3. D) The average MreB polymer number per cell in RodZ cytoplasmic truncation mutants. E) The average MreB polymer number per cell in RodZ periplasmic truncation mutants. D-E) P-value comparisons are made between indicated strain and a strain with full length RodZ (solid red bar). For other comparisons see Table S3. The number above the bars is the number of polymers (B) and the number of cells (C-E) analyzed. Error bars represent 95% confidence intervals. ns P > 0.05, ** P ≤ 0.01, *** P ≤ 0.001

While RodZ does not influence all aspects of MreB polymers, we observed a dramatic decrease in total MreB polymer number per cell (including those <200nm) in the absence of RodZ (WT= 11.6 ± 0.5, *ΔrodZ=* 6.6 ± 0.3) (Fig. 3C). The change in polymer number in Δ*rodZ* cells cannot be attributed to changes in cell shape as mecillinam treatment led to round cells without a concurrent decrease in polymer number (polymer numbers actually increased in these cells, (Fig. S1D). Furthermore, we binned both WT and Δ*rodZ* cells by volume and found that in cells of similar volume Δ*rodZ* had fewer MreB polymers and lacked WT-like geometrically-localized MreB (Fig. S6). Thus, while cell volume does affect polymer number, RodZ increases the number of polymers and promotes curvature localization independently of its effect on cell size and shape.

To further dissect the role of RodZ in assembling MreB polymers, we examined the average polymer number in cells with RodZ truncation mutants. Because MreB curvature localization is more dependent on the cytoplasmic domain of RodZ than its periplasmic domain and MreB binds RodZ in the cytoplasm, we hypothesized that this domain would also control MreB polymer number. As expected, deletion of the cytoplasmic domain of RodZ resulted in a decrease of the number of MreB polymers per cell (Fig. 3D). Surprisingly, we also saw a dramatic reduction in the number of MreB polymers per cell when we truncated the periplasmic domain of RodZ (Fig. 3E). Since the periplasmic domain of RodZ is needed to interact with the cell wall synthesis machinery, these data suggest RodZ could integrate signals from the process of cell growth to feed back on MreB and control polymer number.

We also compared MreB polymer properties in MreB point mutants with or without *rodZ*. We found that specific point mutations altered specific properties of MreB. For example, MreB_E143A_ polymers are longer than WT but have the same MreB polymer angle, while MreB_S14A_ polymers are the same length as WT but the polymer angle is different (Fig. 3A and S4). Interestingly, when comparing MreB_S14A_ in the presence or absence of RodZ, MreB_S14A_ suppressed the RodZ-dependent properties of MreB (curvature localization, polymer number, and membrane-association) (Fig. 3A and S4). MreB_S14A_ was also the strongest suppressor of Δ*rodZ* cell shape, suggesting that MreB_S14A_ functionally restores a majority of the effect of the WT MreB-RodZ interaction. In contrast, MreB_E143A_, a partial suppressor of Δro*dZ* cell shape, suppresses the effects of RodZ on MreB curvature localization but does not suppress the effects of RodZ on MreB polymer number or membrane fraction. MreB_E143A_ also has longer polymers than MreB_WT_ and the length of these polymers increases in the absence of RodZ. Together our results suggest that the different properties of MreB can be modulated independently.

### Cells need multiple, long, and geometrically-localized MreB polymers to grow as uniform rods

Because MreB curvature preference did not always correlate with cylindrical uniformity and MreB parameters can be independently controlled, we sought to determine which properties of MreB best predict cylindrical uniformity. To this end we quantified cell shape and compared MreB properties across a large set of *mreB* and *rodZ* mutants (Supplemental Table 4-mutants and properties). To quantify cylindrical uniformity we relied on our previous analysis^1^ showing that the variation of cell diameter within a single cell (intracellular diameter deviation, IDD) is a quantitative measure of cylindrical uniformity (Fig. S7). We confirmed that the IDD measured from 3D reconstructions also shows a clear separation between cells that are qualitatively classified as uniform rods, irregular rods, and round cells (note that IDD is inversely related to cylindrical uniformity, Fig. S7). We then built a collection of shape comparisons by computing the difference in IDD between two strains (ΔIDD = IDD_strain1_ – IDD_strain2_) (Table 1). Using this nomenclature, a positive ΔIDD describes a comparison where cells of strain 1 are more irregular in their shape than cells of strain 2. For example, Δ*rodZ* cells have a ΔIDD of +0.1 when compared to WT cells (ΔIDD = IDD_ΔrodZ_ – IDD_WT_). We note that in all cases we computed a ΔIDD value that compares two strains with one change, either comparing the same genetic background with or without *rodZ* to assess the impact of RodZ, or comparing WT to different alleles of *mreB* to assess the impact of specific changes to MreB. In addition to ΔIDD, we computed the change in MreB parameters for these same comparisons, choosing as our input a wide variety of scalar quantities (average polymer length, number of polymers, polymer angle, fraction of MreB on the membrane, etc.) and versions of these normalized by the surface area or volume. For the non-scalar metric (curvature localization), we distilled the curvature enrichment profiles into multiple scalars, including the average of the enrichment value for Gaussian curvatures below and above 2 μm^-2^. For a complete list of MreB parameters quantified, see Supplement Table 2.

**Table 1:**
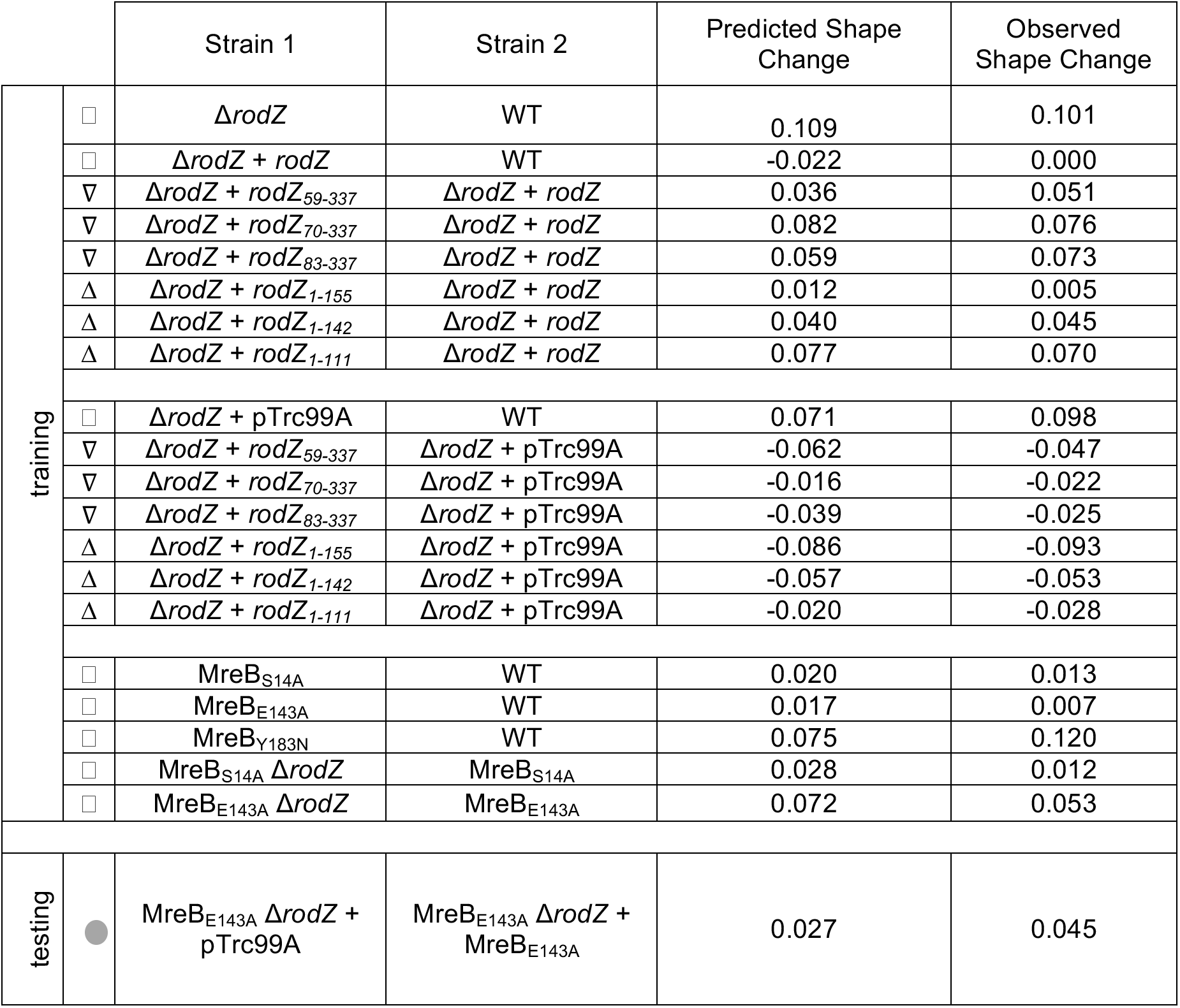
LASSO analysis of MreB’s role in modulating cylindrical uniformity (ΔIDD = IDD_strain1_-IDD_strain2_) A list of strains used to determine LASSO parameters. Predicted change is the change in cell shape (IDD) that we predict from the regression model, while measured shape change is taken from 3D measurements of cells. Δ-RodZ periplasmic truncations, ▽-RodZ cytoplasmic truncations, □-MreB point mutants were strains used to determine the LASSO parameters. Grey circle- comparison used to test the model.

To identify the MreB parameters that were most predictive of changes in cylindrical uniformity we performed a LASSO (Least Absolute Shrinkage and Selection Operator) regression^24^. LASSO is a machine learning method that involves penalizing the absolute size of the regression coefficients. The result is the smallest model within one standard error of the mean of the minimum LASSO regression. Because we did not know *a priori* whether measurements should be normalized per cell, per volume, or per surface area, we used all three normalizations as inputs into the LASSO regression. Combining all of our data, our LASSO analysis resulted in a model with four non-zero terms. These four terms included measures of polymer number, length, and curvature localization with different normalizations. However, a leave-one-out analysis of the data revealed that this was an over fit model (Table S2). To determine which version of normalization was most predictive across different subsampled datasets, we used two different analysis methods (see Materials and Methods). Both methods converged on the same terms: MreB enrichment in regions of low Gaussian curvature (<2 μm^-2^), the total length of MreB polymer in each cell normalized by cell volume, and the number of polymers per cell (Fig. 4, Table S2). The combination of these three parameters was very predictive of the change in cell shape (r^2^=0.93), significantly more than any one parameter alone (r^2^=0.49 (total polymer length), 0.68 (polymer number), and 0.52 (curvature localization)) (Fig 4AB). Importantly, this correlation holds for strain comparisons that have a positive or negative ΔIDD due to truncations in RodZ or MreB point mutations (Fig. 4A).

**Figure 4.**
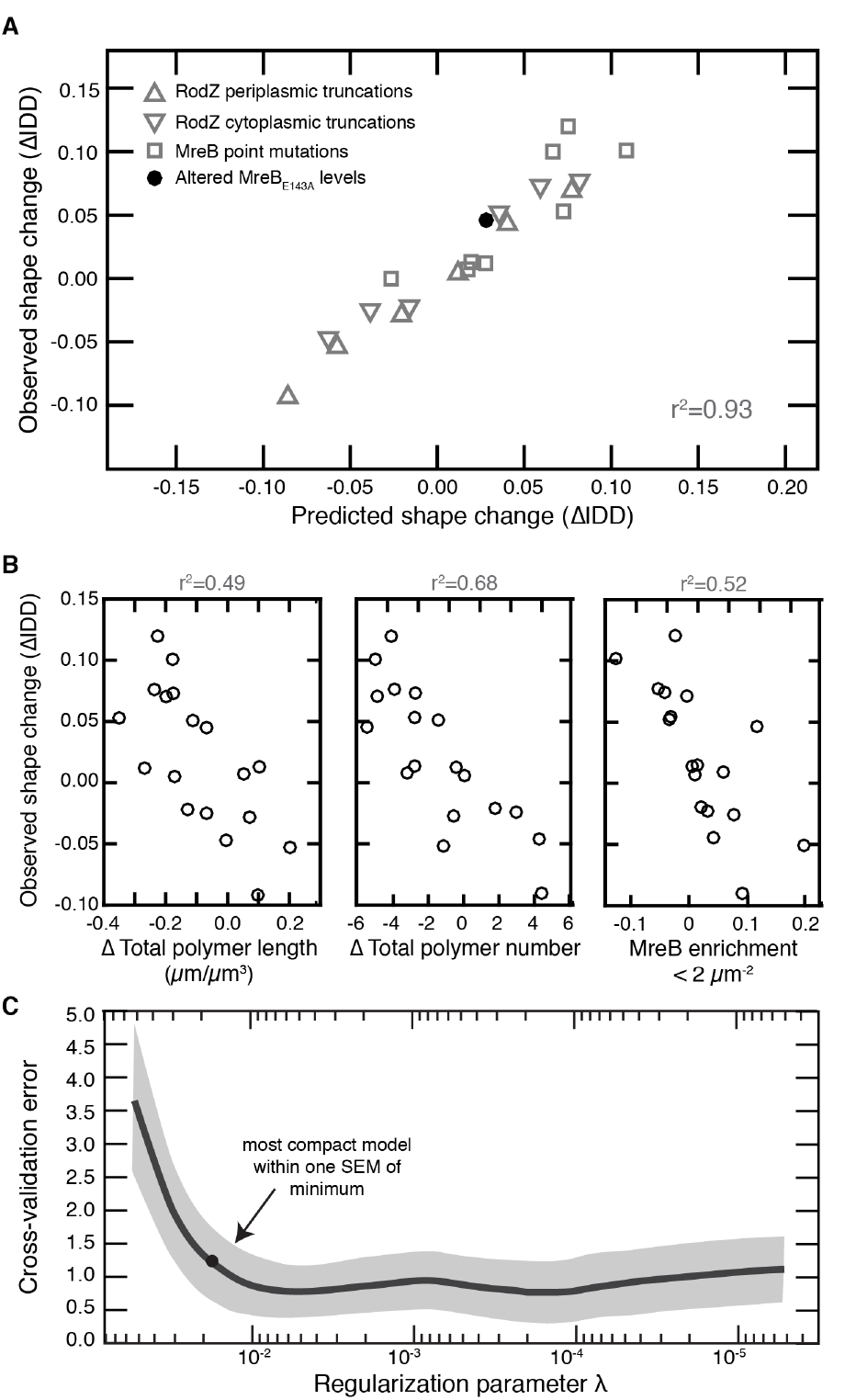
LASSO analysis reveals that rod shape requires many long and geometrically-localized MreB polymers. A) The correlation between observed and predicted cell shape when using the LASSO model combining parameters. See Table 1 for all the strains used and the observed and predicted IDD values. Note that to preserve its use as a test of the model, overexpression of MreB_E143A_ was not used for model selection and training. r^2^ value represents the square of the Pearson correlation coefficient. B) Left-the correlation between observed and predicted cell shape when only using polymer length normalized by volume. Middle-the correlation between observed and predicted cell shape when only using polymer number. Right-the correlation between observed and predicted cell shape when only using average MreB enrichment at Gaussian curvatures below 2 μm^-2^. C) The mean squared error (MSE) of 10-fold cross-validation as a function of the LASSO regularization parameter. The solid curve is the mean MSE and the shaded region represents one standard error of the mean. The dot represents the most compact model within one standard error of the mean from the minimum of the curve. See Table S2 for the coefficients in this model.

The LASSO regression considers the correlations among MreB properties and cell size in order to identify the features of MreB that should be predictive of corresponding changes in cylindrical uniformity (IDD). We thus sought to experimentally test whether the LASSO regression’s result, that changing specific MreB properties, like polymer number, should result in a predictable change in IDD. Because MreB_E143A_ is able to maintain WT-like MreB curvature localization even in the absence of *rodZ*, this particular mutant enabled us to test our hypothesis. Specifically, we found that ectopic expression of MreB_E143A_ in a MreB_E143A_ Δ*rodZ* background increased MreB polymer number (Fig. S8). Importantly, we observed the LASSO-predicted change in shape upon increasing MreB polymer number, as ectopic expression of MreB_E143_ made the cells more rod-like, even though *rodZ* was still absent (Fig. S8, Fig. 4A, Table 1). We also ectopically expressed MreB_WT_ in a MreB_WT_ Δ*rodZ* background, which is not properly curvature-localized. This strain did not restore rod shape, confirming that the MreB_E143_ effect is not a generic consequence of ectopic expression (Fig. S8). These results support our conclusion that MreB-dependent uniform rod shape requires the presence of multiple polymers that are geometrically-localized and collectively long.

## Discussion

Our findings demonstrate that RodZ plays a central role in regulating both MreB’s localization to curved subcellular regions and the number of MreB polymers per cell. This RodZ-mediated regulation also revealed that MreB curvature preference is necessary but not sufficient for cylindrical uniformity, and that cylindrical uniformity requires the presence of multiple long polymers of MreB. Below we discuss the implications of RodZ’s function as an MreB interaction partner. We also present a model in which distinct aspects of MreB control distinct aspects of rod shape determination including rod initiation^2^, centerline curvature^4,7^ (modulated by MreB curvature localization), cell width determination (modulated by MreB angle)^5^, and now, in this current work, cylindrical uniformity (modulated by multiple factors).

### RodZ regulates MreB polymer number, localization, and movement

We previously showed that the transmembrane protein RodZ binds to MreB in the cytoplasm and cell wall synthesis enzymes in the periplasm to couple MreB to cell wall insertion, thereby driving MreB rotation^1^. Here we demonstrate two additional functions for RodZ in regulating MreB curvature preference and polymer number. Using 3D imaging, we show that MreB localizes to areas near zero Gaussian curvature, which causes it to become enriched in the cylindrical region of the cell and avoid the cell poles whose Gaussian curvature is positive. *In vitro* and *in silico* data indicate that MreB polymers have an intrinsic curvature, suggesting that MreB filaments could potentially sense curvatures on their own^9,11,15^. However, our data show that *in vivo, E. coli* MreB_WT_ polymers require RodZ to properly sense cell curvature. Nevertheless, RodZ is not absolutely required for curvature localization as some MreB point mutants can localize in a WT-like pattern even in the absence of RodZ. Understanding the molecular mechanism by which RodZ influences MreB curvature localization will require *in vitro* systems that are currently unavailable. Since RodZ reaches around the MreB polymer and binds it on the side opposite the membrane^20^, we speculate that the binding of RodZ could modulate the intrinsic curvature of MreB^6^. RodZ could also function to rigidify MreB polymers such that the absence of RodZ would cause MreB polymers to become more flexible, allowing them to bend more freely and therefore bind to a wider array of curvatures.

The second new role we discovered for RodZ is in regulating MreB polymer number. Previously, we had attributed changes in polymer number to changes in cell volume^5^. However, mutations in *rodZ* provided a shape-independent way to modulate polymer number, enabling us to conclude that polymer number promotes cylindrical uniformity independently of cell volume. There are several mechanisms by which RodZ could increase MreB polymer number. For example, RodZ could function as a nucleator that stimulates the formation of new polymers, as a severing protein that cuts single polymers into two separate polymers, or as a capping factor that limits polymer growth. We note that a simple polymer stabilization mechanism is unlikely because that would have led to significantly-increased polymer length. Regardless of the mechanistic details, whose dissection will require future *in vitro* studies, our findings represent the first identification of a factor that enhances MreB polymer formation.

Interestingly, our RodZ truncation and MreB mutant analyses suggest that the functions of RodZ in promoting MreB rotation, polymer formation, and geometric localization are genetically separable. Thus, RodZ appears to use its cytoplasmic and periplasmic domains to coordinate multiple aspects of MreB, including acting upstream of MreB assembly to regulate its polymer properties and downstream of MreB assembly to regulate its coupling to the movement of the cell wall synthesis machinery. Such modularity in a transmembrane protein is an appealing way for the cell to tune the properties of MreB, perhaps enabling optimization of MreB in response to different growth conditions.

### Cylindrical uniformity requires multiple long and curvature-localized polymers

In previous studies, our lab and others determined that MreB mediates multiple aspects of rod shape determination^1,2,4,5^. MreB localization helps straighten rods and initiate new rods out of spheres, while MreB angle is correlated with the average width of a rod^2,5^, even in the absence of RodZ (Fig. S9). Here we show that MreB also determines the cylindrical uniformity of rods (Fig. 5). Specifically, a machine learning analysis (LASSO) of the correlations between MreB properties and rod shape revealed that a combination of modulating polymer number and total polymer length, along with the correct curvature localization is sufficient to accurately predict rod shape changes. While the LASSO analysis cannot distinguish whether MreB polymer number and total polymeric content (sum of the length of all polymers) directly impact cell shape or are correlates of other cellular processes, our result supports previous studies suggesting that MreB forms multiple independent structures distributed throughout the cylindrical portion of the cell^1,5,12,13,25,26^. Because both polymer number and total polymeric content are important, we predict neither one long polymer nor a multitude of small polymers are sufficient to generate cylindrically uniform cells. MreB senses local cell curvature and directs cell wall synthesis to those sites, which in turn locally changes cell shape. Thus, our data suggest that maintaining a rod shape with uniform diameter requires multiple MreB structures to make enough local shape measurements to direct the overall emergence of rod shape. In addition to promoting curvature sensing, long MreB structures could also distribute the area along which new cell wall material is being inserted^4,14,27^.

**Figure 5.**
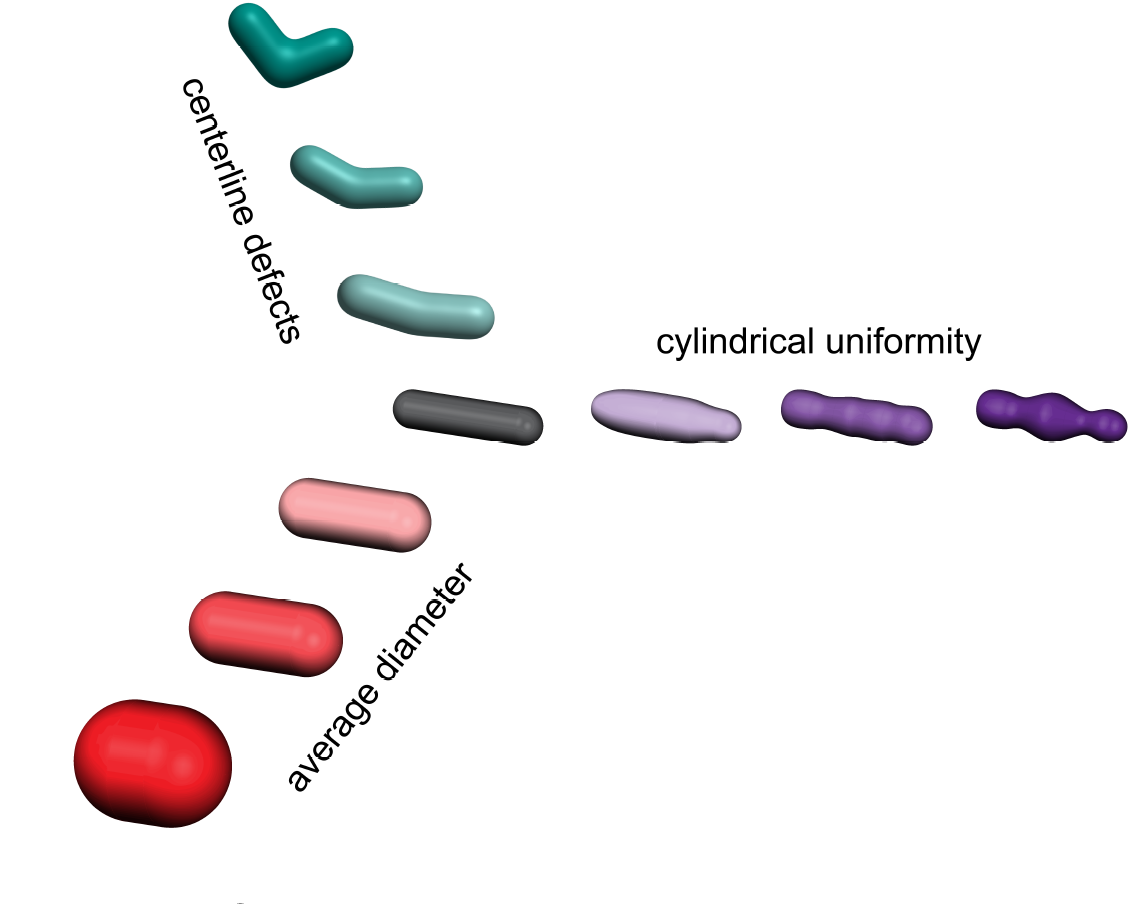
Simple straight rod like cell shapes require multiple parameters to describe them. A straight rod is defined by its centerline curvature, cylindrical uniformity, and diameter. Deviations in any of these properties result in nonstraight rods in extreme cases, and qualitatively ambiguous rods when only small changes occur. Teal-as centerline defects increase in magnitude cells become more bent until they are no longer straight. Purple-as cylindrical uniformity decreases cells exhibit increase fluctuations in their diameter along the long axis. Red-as width increases cells become more sphere like and less rod like. For each of these shape descriptors, a quantitative metric of shape provides a continuous rather than a binary description of rod vs non-rod.

Our model for cylindrical uniformity predicts changes in cell shape (ΔIDD) in a variety of backgrounds (mutations in both *mreB* and *rodZ)*, and thus represents an additive model of cell shape. The fact that the aspects of MreB that predict cylindrical uniformity (polymer number, length, and curvature localization) are distinct from the aspect of MreB that predicts cell width (polymer angle) suggests that the absolute shape of the cell (width, straightness, uniformity, etc.) is a complicated function where different cell shape properties can be tuned independently (Fig. 5). Thus, while MreB is emerging as a central coordinator of rod shape, there are other components downstream of MreB necessary to physically build the cell wall that are important for determining rod shape. Because RodZ influences both polymer number and MreB curvature localization, it will be important for future studies to unravel the specific contributions of curvature localization to the various aspects of rod shape formation.

